# “*Staphylococcus aureus* SigS induces expression of a regulatory protein pair that modulate its mRNA stability”

**DOI:** 10.1101/2022.10.19.512975

**Authors:** Amer Al Ali, Jamilah Alsulami, Joseph I Aubee, Ayotimofe Idowu, Brooke R. Tomlinson, Emily A. Felton, Jessica K. Jackson, Lindsey Shaw, Karl M Thompson

## Abstract

SigS is the sole extracytoplasmic function sigma in *S. aureus* and is necessary for virulence, immune evasion, as well as surviving exposure to toxic chemicals and environmental stressors. Despite the contribution of SigS to a myriad of critical phenotypes, the downstream effectors of the SigS-dependent *S. aureus* pathogenesis, immune evasion, and stress response remain elusive. To address this knowledge gap, we analyzed the *S. aureus* transcriptome following transient over-expression of SigS. We identified a bi-cistronic transcript, up-regulated by 1000-fold, containing two mid-sized genes each containing single domains of unknown function (DUF). We renamed these genes *sroA* (<u>S</u>igS <u>r</u>egulated *<u>o</u>rf<u>A</u>*) and *sroB* (<u>S</u>igS <u>r</u>egulated *<u>o</u>rf<u>B</u>*). We demonstrated that the SigS regulation of the *sroAB* operon is direct using in vitro transcription analysis. Using northern blot analysis, we also demonstrated that SroA and SroB have opposing auto- regulatory functions on the transcriptional architecture of the *sigS* locus; with SroA stimulated SigS mRNA levels and SroB stimulating s750 (SigS antisense) levels. We hypothesized that these this opposing regulatory effects were due to a direct interaction. We demonstrated an interaction between SroA and SroB using an in-vivo surrogate genetics approach via Bacterial Two Hybrid. We demonstrated that the SroA effect on SigS is at the post-transcriptional level of mRNA stability, highlighting a mechanism likely used by *S. aureus* to tightly control SigS levels. Finally, we demonstrate that the *sroAB* locus promotes virulence in a female murine pneumonia model of infection.

## INTRODUCTION

*Staphylococcus aureus* is a member of the firmicute (low G-C% genome gram-positive) phylum of bacteria and is further distinguished by its yellow pigment (via Staphyloxanthin), spherical shape (coccus), and absence of spore forming ability. *S. aureus* is a formidable human pathogen capable of causing dangerous infections. In 2017, there were approximately 120,000 *S. aureus* bloodstream infections nationwide, resulting in 20,000 fatalities (1). *S. aureus* can cause respiratory co-infections in patients suffering from SARS-COV-2 (COVID-19) or Influenza, resulting in a COVID or Flu / necrotizing pneumonia co-morbidity (2, 3). This is further exacerbated by the existence of multi-drug resistant clinical isolates, most notably Methicillin Resistant *S. aureus* (MRSA) (4). MRSA infections are generally treated with last line antibiotics such as Vancomycin and Daptomycin (5). Unfortunately, *S. aureus* isolates with increased resistance to both Vancomycin (Vancomycin Intermediate *S. aureus* – VISA) and Daptomycin (Dap^R^) have been identified (5, 6). The incidence of antimicrobial resistance has increased as a result of the COVID-19 pandemic (7).

*S. aureus* virulence is facilitated by a large repertoire of virulence factors that facilitate the pathogenesis process (8). Virulence factor gene regulation is critical for *S. aureus* stress adaptation and evasion of the host response, both of which are necessary for the ability of *S. aureus* to establish a productive infection (8). *S. aureus* virulence gene regulators include Staphylococcal accessory regulator (Sar) proteins, RNAIII, and a host of two component signal transduction systems (8). In addition, alternative sigma factors can regulate virulence gene expression in *S. aureus*. *S. aureus* has three alternative sigma factors: SigB, SigH, and SigS (9–11).

SigS is the sole *S. aureus* extracytoplasmic function (ECF) sigma factor and is necessary for stress adaptation and immune evasion (11, 12). In the absence of sigS, S. aureus is attenuated for virulence and defective in producing joint inflammation in a mouse model of septic arthritis (11). *S. aureus* Long term cell viability in vitro and survival during extreme heat shock (>55°C) both require SigS (11). Finally, *S. aureus* sigS mutants have increased sensitivity to cell wall targeting antibiotics, detergents, DNA damaging agents, and the innate immune system components (11, 13). Yet, the mechanism whereby SigS senses these stressors and promotes adaptive phenotypes is not clear. There appears to be some continuity between adaptive phenotypes and the induction of *sigS* expression as exposure DNA damaging agent MMS, H_2_O_2_, NaOH, immune cell components, and macrophage phagocytosis stimulate expression of SigS (13). However, the regulation of SigS is complex as basal expression is low in the absence of inducing signals and it has 4 promoters controlling its transcriptional initiation (11, 13, 14). Genetic and biochemical screens identified CymR and KdpE as direct regulators, and ArlR and LacR as indirect regulators, of *sigS* transcription (15). Prior to this work, little was known about the post-transcriptional regulation of *sigS* expression or the direct targets of SigS that could promote virulence and stress adaptation.

ECF sigma factors usually mediate their stress response through protein or small RNA mediators that are their direct targets (16–20). In *E. coli*, and other enteric bacteria, the ECF sigma factor σ^E^ has a relatively large regulon composed of periplasmic proteases and small RNAs that restore envelope homeostasis through the fine tuning of outer membrane protein levels (16–20). In *B. subtilis* and other firmicutes, ECF sigma factors also have relatively large regulons that ensure envelope homeostasis in the presence of cell wall stressors (21, 22). Based on these observations in other bacterial systems, we hypothesized that the SigS would mediate its stress adaptation, virulence, and immune evasion through the action of such direct regulatory targets.

In this work, we analyzed the *S. aureus* transcriptome following transient *sigS* over- expression. Consequently, we identified a major SigS effector locus with a previously uncharacterized regulatory protein pair tandemly encoded on a bis-cistronic transcript (encoding SAOUHSC_00622 and an upstream ORF). Each of these ORFs are comprised of a single domain of unknown function (DUF): DUF1659 and DUF2922, respectively. We renamed these genes *sroAB* (<u>S</u>igS <u>r</u>egulated *<u>o</u>rf<u>A</u>* and *orf<u>B</u>*). Further, we show that SroA and SroB have opposing regulatory effects on the *sigS* locus. We also uncovered positive feedback regulation as SroA acts to stabilize the *sigS* transcript, thereby promoting SigS accumulation in the absence of SroB. Finally, we demonstrate that the *sroAB* locus is necessary for full virulence in a murine model of pneumonia. This work provides insight into a network of tight controlled of the cryptic SigS sigma factor in *S. aureus*.

## MATERIALS AND METHODS

### Media and Growth Conditions

*E. coli* strains used for propagation of recombinant DNA were grown in Luria Bertani Lennox broth (LB), or LB agar plates, supplemented with ampicillin to a final concentration of 100 μg / mL (LB-amp), kanamycin to a final concentration of 25 μg / mL (LB-kan), or chloramphenicol to a final concentration of 10 μg / mL or 25 μg / mL (LB-Cm). *E. coli* strains co-electroporated with Bacterial Two-Hybrid (BTH) clones were grown in LB agar plates supplemented with ampicillin to a final concentration of 100 μg / mL and kanamycin to a final concentration of 25 μg / mL (LB-Amp-Kan). Overnight cultures of *E. coli* were grown in 5 mL of LB within a roller drum placed inside of a microbiological incubator at 30°C or 37°C. All *S. aureus* strains were grown in Tryptic Soy Broth (TSB). Overnight cultures of *S. aureus* were grown in 5 mL of TSB within a roller drum placed inside of a microbiological incubator at 37°C. *S. aureus* strains containing pEPSA5 or its derivatives were grown in TSB supplemented with Chloramphenicol to a final concentration of 10 μg / mL (TSB-Cm). Phage 80 lysates were created, using the top agar method, on *S. aureus* RN4220 cells grown on TSA-Cm, supplemented with CaCl_2_ to a final concentration of 5mM. *S. aureus* transformants were grown on TSB-Cm plates (TSA-Cm) and transductants were selected on TSA-Cm supplemented with 5 mM Sodium Citrate (NaCi).

### Strains, Plasmids, and Oligonucleotides

All strains and plasmids are listed in Table 1. *Staphylococcus aureus* cells used in this study were derivatives of NCTC8325 (Table 1) (8). *S. aureus* plasmid expression vectors were electroporated in restriction minus *S. aureus* strain RN4220 as previously described (23, 24). Plasmids, and previously constructed *S. aureus* mutations, were transferred from RN4220 or other *S. aureus* strains into SH1000 by transduction using ϕ80α as previously described (25). Mutations constructed in RN4220 or SH1000 using pIMAY* as previously described (26). The *E. coli* strain used in this study for cloning purposes was NEB-5α (New England Biolabs) (Table 1). Expression plasmids used in this study are listed in Table 2 and are derivatives of the pEPSA5, an *E. coli* / *S. aureus* shuttle vector containing a xylose inducible promoter and its cognate regulator XylR (Table 2). Oligonucleotides used for PCR amplification or as biotinylated Northern Blot probes are listed in Table 3. Plasmids used for Bacterial Two Hybrid Analysis were derivatives of pEB355 or pEB354 (27, 28). *Escherichia coli* NEB- 5α (New England Biolabs) cells were used for all cloning reactions. Clones containing inserts or site-directed point mutants, as described below, were all identified by colony PCR and verified by PCR amplification and DNA sequencing of the purified recombinant plasmids.

**Table 1.**
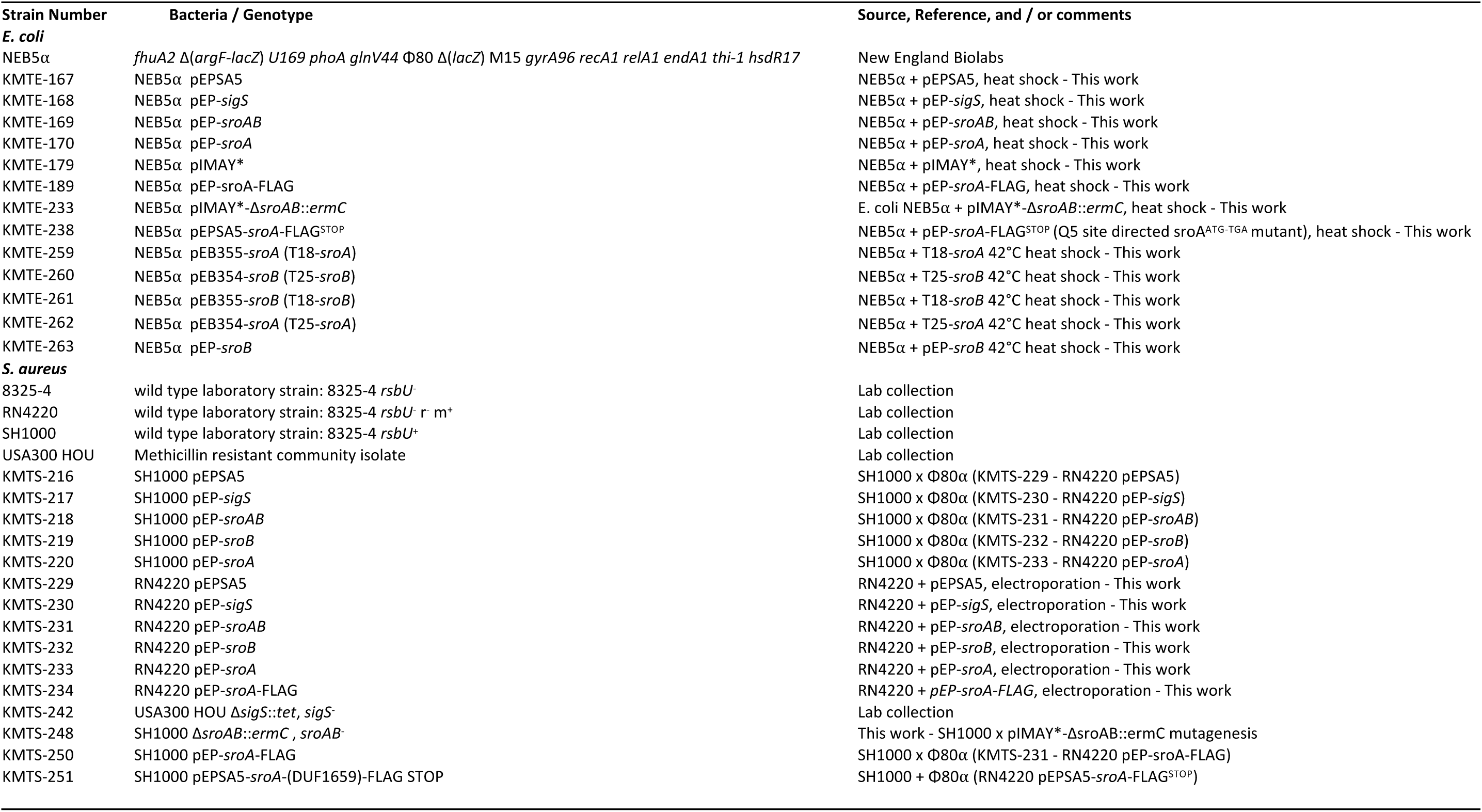
Strain List.

**Table 2.**
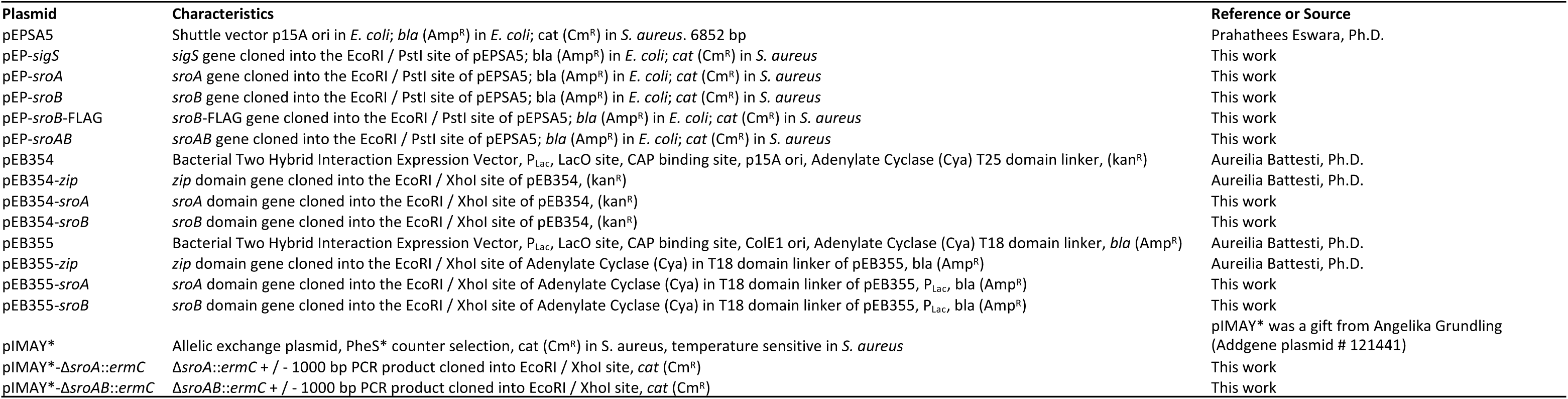
Plasmid List.

**Table 3.**
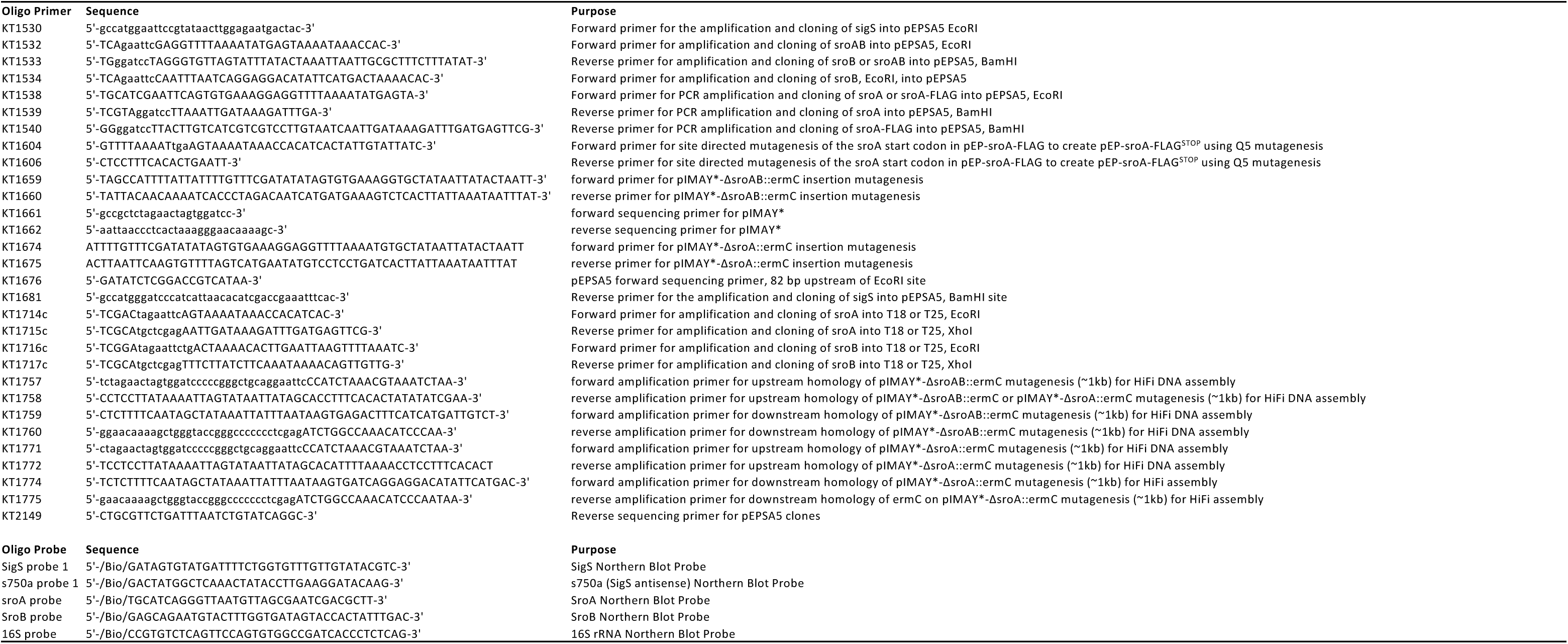
Synthetic Oligonucleotides used as primers or probes.

#### Construction of pEPSA5 derivatives

For our studies in this work, we created several plasmid-based xylose-inducible constructs via standard recombinant DNA technology techniques by cloning desired genes for expression into pEPSA5. We created pEPSA5 derivatives containing *sigS*, *sroA*, *sroB*, *sroAB*, and *sroA*-FLAG. Briefly, we amplified the respective genes from the SH1000 genomic DNA using their corresponding primers oligonucleotide primers as listed in Table 3. Both purified pEPSA5 and the PCR products to be inserted were digested with EcoRI and BamHI and ligated using instant sticky-end ligase. Ligation reactions were transformed into *E. coli* NEB-5α (New England Biolabs) cells and screened by colony PCR. To construct a nonsense mutation in the *sroA* start codon, within the pEP-*sroA* construct (pEP-*sroA*-FLAG^STOP^), we executed mutagenic PCR oligonucleotide primers KT1604 and KT1606 and the Q5^®^ Site-Directed Mutagenesis Kit (New England Biolabs) according to manufacturer’s instructions.

#### Construction of pIMAY* derived mutagenesis vectors for *sroA* and *sroAB* mutagenesis

We constructed recombinant plasmids containing allelic exchange substrates for the creation of erythromycin-marked deletion insertion mutations of *sroA* or *sroAB*, (pIMAY*-!ι*sroA*::*ermC* and pIMAY*-!ι*sroAB*::*ermC*, respectively). We created pIMAY*-!ι*sroAB*::*ermC* using HiFi DNA Assembly. First, we then amplified three different DNA sequences: 1.) “!ι*sroAB*::*ermC* up” from *S. aureus* SH1000 genomic DNA using oligonucleotide primers KT1757 and KT1758; 2.) “!ι*sroAB*::*ermC* internal” from *S. aureus* shuttle vector pCN51 using oligonucleotide primers KT1659 and KT1660; and 3.) “!ι*sroAB::ermC* down” from *S. aureus* SH1000 genomic DNA using oligonucleotide primers KT1759 and KT1760. The “!ι*sroAB*::*ermC* up” PCR product has approximately 30 bp of homology to the region upstream of the multiple cloning site (MCS) of pIMAY*, on its 5’ end, and approximately 30 bp of homology to the 5’ end of the “!ι*sroAB*::*ermC* internal” PCR product, on its 3’ end. The “!ι*sroAB*::*ermC* down” PCR product has approximately 30 bp of homology to the region downstream of the multiple cloning site (MCS) of pIMAY*, on its 3’ end, and approximately 30 bp of homology to the 3’ end of the “!ι*sroAB*::*ermC* internal” PCR product, on its 5’ end. We then digested pIMAY*-!ι*sroAB*::*ermC*, we digested pIMAY* with both EcoRI and XhoI. The resulting pIMAY* EcoRI / XhoI, and the three PCR products were joined using the NEBuilder^®^ HiFi DNA Assembly Master Mix (New England Biolabs) according to manufacturer’s recommendations and transformed into NEB5α cells (New England Biolabs). Transformants were selected on LB-Cm plates. We created pIMAY*-!ι*sroA*::*ermC* using an identical strategy. 1.) “!ι*sroA*::*ermC* up” from *S. aureus* SH1000 genomic DNA using oligonucleotide primers KT1771 and KT1772; 2.) “!ι*sroA*::*ermC* internal” from *S. aureus* shuttle vector pCN51 using oligonucleotide primers KT1674 and KT1675; and 3.) “!ι*sroA*::*ermC* down” from *S. aureus* SH1000 genomic DNA using oligonucleotide primers KT1774 and KT1775. The subsequent steps were identical to those described above used to construct pIMAY*-!ι*sroAB*::*ermC*.

#### Construction of plasmids for Bacterial Two Hybrid (BTH) Analysis

We created recombinant plasmids containing monomers of adenylate cyclase cloned in frame with the *sroA* or *sroB* genes into BTH vectors pEB355 or pEB354 (27, 28). The *sroA* and *sroB* genes were amplified by PCR, using their corresponding oligonucleotide primers listed in Table 3 and digested with EcoRI and BamHI. We then ligated the purified *sroA* or *sroB* PCR product digests with the purified pEB355 or pEB354 EcoRI / BamHI digests using the Instant Sticky End Ligase (New England Biolabs) and transformed them into chemically competent NEB-5α cells (New England Biolabs) according to manufacturers’ instructions. Transformants were selected on LB-Amp plates.

### Total RNA Isolation

Total RNA was isolated using the Fast RNA Pro^TM^ Blue Kit (MP Biomedicals). Briefly, *S. aureus* cells of interest were harvested from 5-50 mL of culture at the selected condition (OD_600_ and or specific experimental treatment). The pellets were resuspended in 1 mL of RNA Pro Solution, added to silica beads, and homogenized using the Precellys® 24 Dual Homogenizer (Bertin Corporation) with the Cryolys for Precellys® adaptor to ensure integrity of RNA following heat exposure. Following lysis, RNA extraction was further executed via Chloroform extraction and Ethanol precipitation overnight at -80°C. RNA cell pellets were resuspended in 50-100 μL of RNase-free H_2_O. RNA concentration was measured using the Nanodrop.

### RNA Sequencing

RNA sequencing and data analysis was performed as described previously (29). Briefly, the SH1000 wild-type carrying either empty pEPSA5 or pEP-*sigS* was grown overnight as detailed above. Next, these cultures were used to inoculate fresh TSB, and allowed to grow for a further 3h. After this time, these cultures were used to seed fresh TSB at an OD_600_ of 0.05. Strains were allowed to grow until exponential phase (OD_600_ = 0.3) before the addition of xylose to a final concentration of 2%, to induce expression of *sigS*. These cultures were grown for 30 mins before samples were combined with 5mL ice-cold PBS and subject to centrifugation. Total RNA extractions were performed using a Qiagen RNAeasy Kit and DNA was removed with the Ambion TURBO DNA-free kit. RNA quality was determined using an Agilent 2100 Bioanalyzer with an RNA 6000 Nano kit to confirm RNA integrity (RIN). Only samples with a RIN of 9.7 or higher were used in this study. Triplicate samples for each strain, from independently grown cultures, were then pooled at equal RNA concentrations, followed by rRNA removal using a MICROBExpress Bacterial mRNA enrichment kit. Efficiency of rRNA removal was confirmed using an Agilent 2100 Bioanalyzer with an RNA 6000 Nano kit. These mRNA samples were then subject library preparation using a Truseq Stranded mRNA Kit (Illumina) with the mRNA enrichment steps omitted. Fragment size, quantity and quality were assessed using an Agilent 2100 Bioanalyzer with an RNA 6000 Nano kit. Library concentrations for the pooling of barcoded samples was assessed with a KAPA Library Quantification kit. Samples were run on an Illumina NextSeq with a 150- cycle NextSeq Mid Output Kit v2.5. Experimental data from this study were deposited in the NCBI Gene Expression Omnibus (GEO) database (GEO accession number GSE215075). Data was exported from BaseSpace (Illumina) in fastq format and uploaded to CLC Genomics Workbench for analysis. Data was aligned to the NCTC 8325 reference genome (NC_007795.1), and comparisons were carried out following quantile normalization via the Qiagen Bioinformatics experimental fold change feature.

### In vitro transcription of *sroA*

One unit of RNA polymerase Core Enzyme (New England Biolabs) was added reconstituted with 1 μg of purified recombinant σ^S^ (vendor) in 1 μL of 5X RNA Polymerase Reaction Buffer (New England Biolabs) and incubated at 4°C for 15 minutes. After incubation, 1 μg of a purified PCR product corresponding to the *sroA* coding region and 980 bp of the *sroA* upstream region, region was added to the σ^S^-core--RNA polymerase mixture and incubated at 37°C for 15 minutes. Then, rNTPs were added to initiate transcription and the reaction was incubated at 37°C for 30 minutes. After this incubation, the RNA was purified twice by acid phenol chloroform extraction followed by ethanol precipitation. RNA was also purified by TURBO DNA-free^TM^ Kit (Thermo Scientific) following the manufacturer’s protocol. 1-step RT-PCR reaction then was executed on the purified RNA using two primers KT1578 / KT1579 (creating a product corresponding to the start codon of *sroA* to the termination codon of *sroA*). This experiment was repeated, with two controls, purified σ^S^ and core-RNA polymerase. RT-PCR reactions were then run on a 2% agarose gel and the image was obtained using the FluorChem R system.

### Bacterial Two Hybrid Analysis

We analyzed protein-protein interactions in vivo using a previously described Bacterial Two Hybrid System (27, 28). Briefly we co-electroporated putative protein interaction pairs (pEB355-*sroA* and pEB354-*sroB* or pEB355-*sroB* and pEB354-*sroA*), the empty vector control pairs (pEB355 and pEB354), or the positive control pairs (pEB355-*zip* and pEB354-*zip*) into *E. coli* BTH101. We allowed the cells to recover at 30°C for several hours and plated them on LB agar plates supplemented with ampicillin and kanamycin. Then, several colonies were inoculated into 5 mL of LB-amp- *kan* media supplemented with IPTG to a final concentration of 100 μM and inoculated overnight at 30°C. A 10 mL aliquot of the overnight cultures was spotted onto MacConkey-Maltose plates supplemented with IPTG to a final concentration of 100 mM and incubated at 30°C for 24-48 hours to measure the Lac phenotype. Alternatively, a 100 mL aliquot of the overnight culture was subjected to a kinetic β-galactosidase assay.

### SigS Rifampicin Chase Assay

We grew SH1000 containing either pEPSA5 (empty vector control) or pEP-*sroA* in TSB-Cm supplemented with MMS to a final concentration of 25 mM, at 37°C to an OD_600_ of 0.5. We harvested the cells to remove supplemental MMS and resuspended cells in fresh TSB-Cm supplemented with Xylose to a final concentration of 2% and incubated the resuspended cultures for an additional 30 minutes. We then harvested the cells by centrifugation to remove supplemental Xylose and resuspended the cells in fresh TSB-Cm. We then added Rifampicin to a final concentration of 500 mM, to stop intracellular transcription, and isolated total RNA at 2– 5-minute intervals following Rifampicin supplementation. We then subjected these RNAs to Northern Blot analysis.

### Northern Blot Analysis

A 1% MOPS Agarose gel was used for resolution of total RNA. The agarose gel and buffer were created using UltraPure^TM^ Agarose (Invitrogen^TM^) in 1X MOPS Buffer, diluted 1:10 from 10X MOPS Buffer (Quality Biological, Inc) mixed with DEPC water. The gel was pre-run at 100v for 40-minutes. Then, 2-5ug of total RNA was mixed with 2X volume of loading buffer (500ul Formamide, 100ul 10X MOPS, 100ul (80% glycerol 0.2% bromophenol blue) and 120ul Formaldehyde, 2ul (10mg/ml EtBr). The samples were then heated at 65° C for 15-minutes and loaded onto the gel for fractionation by gel electrophoresis for 40-minutes at 100v. After the completion of the fractionation, the gel was soaked in 0.05 M NaOH solution for 20-minutes, and then upside down in 20X SSC solution for 1-hour. The membrane was transferred using the capillary method and crosslinked with short-wave UV light to fix the transferred RNA onto the membrane. The crosslinked membrane was then pre-hybridized with 5ml of PerfectHyb^TM^ Plus Hybridization Buffer (Millipore Sigma) for 2-hours. Then a biotinylated DNA probe corresponding to either SigS or 16S rRNA was added to the Hybridization Buffer to a final concentration of 500 ng / mL and hybridized at 42° C for 2 hours. The membranes were then processed using stringency washes and developed using the Chemiluminescent Detection Kit according to manufacturer’s recommendations. The chemiluminescent signal was detected using the Fluorochem R (Protein Simple). Northern blot densitometry signals were quantified using Image J and analyzed using Graphpad Prism.

### Total Protein Isolation

To isolate total protein, *S. aureus* cells were harvested from a 10mL culture aliquot. The pellet was resuspended in 1mL of 1X PBS. The cell-PBS suspension was processed twice in the Precellys Dual Homogenizer for 40 seconds at a setting of 5000 and centrifuged at 15,000 g for 5 minutes at 4°C. The proteins were precipitated from the solution via treatment with Trichloroacetic acid (TCA) to a final concentration of 25% on ice followed by centrifugation at 15,000 rpm for 5 minutes at 4°C. The pellet was washed using 200 μL of 100% ice cold acetone (Thermo Scientific). The pellet was resuspended in 50 μL of 1X PBS and protein concentration was using The DC Protein Assay Kit (Biorad).

### SDS-PAGE and Western Blot Analysis

30 μg of total protein mixed with 4X NuPAGE LDS Sample Buffer (Thermo Scientific) and NuPAGE Sample Reducing Agent (Thermo Scientific) heated at 70oC for 10 minutes and then cooled on ice for 2 minutes. Then the sample was loaded onto a 12% Bis-Tris Mini-protein Gel (Thermo Scientific) and subjected to electrophoresis using 1X MES Buffer (Thermo Scientific) at 100 volts for 1 hour. The protein gel was then transferred to a 0.45 μm pore-sized nitrocellulose membrane using the Trans-Blot® TurboTM System (Biorad), according to manufacturer’s instructions. The membrane was washed for five minutes, three times, with PBST Buffer. Then, the membrane was washed in Blocking solution (5% Blotting-Grade nonfat milk in 1X PBST Buffer) for 30 to 120 minutes at room temperature with gentile agitation. The membrane was then washed three times with 1X PBS-T with gentle agitation. Next, the membrane was incubated in the Blocking solution containing the primary antibody to the FLAG epitope tag, α-FLAG (Sigma-Adrich), diluted 1:10,000 overnight. Following three washes with PBST for five minutes. Then the membrane was incubated for 2 hours in Blocking solution containing the secondary antibody diluted 1:25,000. The membrane was then washed three times with PBST for five minutes. The membrane was then incubated with Chemiluminescent Detection Solution (Thermo Scientific), prepared by mixing 1 ml of Chemiluminescent substrate and 50 μl of Substrate Enhancer for 5 minutes. The signal was developed using the FluorChem R System (Protein Simple).

### Mouse pneumonia model of infection

Male and female 6-week-old C57BL/6J mice were purchased from Charles River Laboratories and allowed to acclimate for 1 week prior to infection. To prepare inocula, cultures of the wild-type and mutant strains were grown overnight (37°C, 250 rpm), before being diluted 1:100 in fresh TSB. After this time, strains were grown for 3 h, before being standardized to an OD_600_ of 0.05. These cultures were then allowed to grow for 15 hours before the CFU/mL was determined. This process was repeated three separate times, and the average CFU/ml was used to calculate the volume of bacteria required to generate a 10 ml inoculum of 1 x 10^8^ CFU/30 µl. On the day of infection, cultures were grown as described and the calculated volume of bacteria was harvested by centrifugation, washed with PBS, before being resuspended in 10 ml PBS. Mice were anesthetized with isoflurane and then inoculated intranasally with 30 µl of the prepared bacteria suspension. Infections were monitored for 24 hours or until mice reached a premoribund state, at which point they were euthanized. Lungs were harvested and homogenized in 2 ml PBS, and the bacterial burden (CFU/mL) was determined by serial dilution and plating. A Mann-Whitney test was used to determine statistical significance between the mutant and wild-type infected mice.

### Statistical Analysis

All statistical analysis was executed using Prism 9 software (Graphpad).

## RESULTS

### *S. aureus* transcriptome architecture following SigS over-expression

To identify downstream regulators of *sigS* that may promote virulence, immune evasion, or antibiotic resistance, we analyzed the *S. aureus* transcriptome following SigS over-expression (Figure 1A). Briefly, exponentially growing S. aureus cultures were induced for 30 mins before RNA isolation and RNA sequencing (Figure 1A). Approximately 50 genes were up-regulated, and 75 genes were down-regulated in *S. aureus* SH1000 by at least two-fold following transient over-expression of SigS (Figure 1B). Interestingly, many of the SigS up-regulated transcripts were those involved in immune evasion: *spa* (Staphylococcal Protein A), *hlgC* (Leukocidin S Subunit), *hlgB* (Leukocidin F Subunit), *sbi* (IgG binding protein), *lukA* (gamma-hemolysin subunit), *lukB* (gamma-hemolysin subunit B), and *hmp* (nitric oxide dioxygenase) (Figure 1C). Three small RNAs were moderately induced (2-5-fold) following SigS induction: Ssr54, JKD6008sRNA037, JKD6008sRNA401, and JKD6008sRNA140 (Figure 1D).

**Figure 1.**
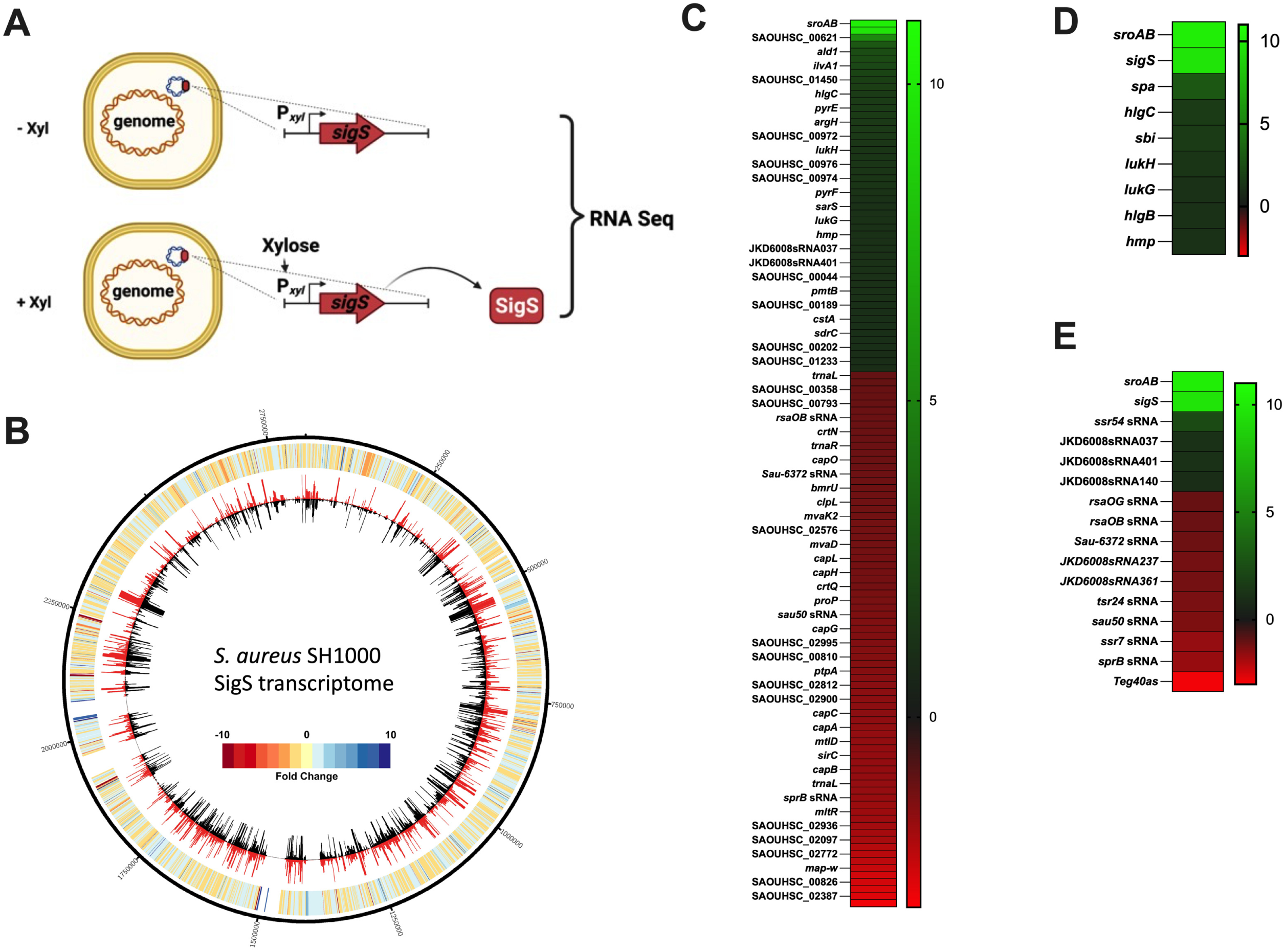
*S. aureus* SH1000 transcriptome following SigS over-expression. A.) Genetic constructs and experimental design for SigS transcriptome experiment. Total RNA was isolated from *S. aureus* strains carrying plasmid-based xylose-inducible sigS construct, or empty vector control, was grown in the presence of xylose to induce over- expression of SigS. B.) CIRCOS file demonstrating *S. aureus* SH1000 transcriptome following over-expression of SigS C.) Heat map of RNA sequencing data depicting *S. aureus* SH1000 transcripts whose expression was modulated by at least 2-fold throughout the transcriptome following SigS induction. D.) Heat map of SigS modulated transcripts involved in immune evasion that were induced by at least 2-fold in reference to SigS and SroAB transcript levels. E.) Heat map of SigS modulated transcripts for small RNAs modulated by at least 2-fold in reference to SigS and SroA transcript levels.

There were 75 genes whose expression was downregulated in response to transient induction of SigS (Figure 1B). Many of the SigS down-regulated genes included genes of unknown function with no functional annotation (Figure 1B). Fifty-three of these SigS down-regulated genes were part of the regulon of the alternative sigma factor SigB (Table S1). Thirteen of the SigS repressed genes that were also a part of the SigB regulon, are also repressed by CodY (Table S1). Interestingly, the *capABCDEFGHJKLMN* operon involved in capsule biosynthesis was repressed, which contrasts with other immune evasion genes noted above that were stimulated in response to induction of SigS. The expression of eight small RNAs was also decreased by at least two-fold following SigS overexpression: *rsaOG*, *rsaOB*, *sau-6372*, *tsr24*, *sau50*, *ssr7*, *sprB*, *teg40as*. Two of these, SprB and Teg40as, are activated by SigB (Table S1 and Figure 1D).

### A novel regulatory protein pair with unknown function are strongly induced by SigS

Beyond the examples noted above, the most highly upregulated genes that emerged from our SigS transcriptome experiment were those that specify two relatively short uncharacterized proteins. These are encoded in tandem, on a single polycistronic transcript, and were induced 1000-fold upon transient over-expression of SigS (Figure 1). Herein we rename them <u>S</u>igS <u>R</u>egulated <u>O</u>RF <u>A</u> (SroA) and <u>S</u>igS <u>R</u>egulated <u>O</u>RF <u>B</u> (SroB). The *sroA* gene encodes a 67 amino acid protein with a DUF1659 domain (Figures 2A). The *sroB* gene encodes a 74 amino acid protein with a DUF2922 domain (Figure 2A). There is a 16 bp region in between the stop codon of *sroA* and the start codon of *sroB* that contains a canonical Shine Dalgarno sequence (AGGAGG) that is presumably a ribosome binding site (Figure 2). There is also a canonical Shine Dalgarno sequence (AGGAGG), starting 16 bp upstream of the predicted *sroA* start codon. While the DUF1659 and DUF2922 domains are widely present in many bacterial species, there is little to no information about their function. Importantly, the amino acid sequences of the *S. aureus* SroA and SroB proteins are conserved only within the Staphylococcal family. Computational analysis of SroA and SroB amino acid sequences using the web-based String database (https://string-db.org/), demonstrate strict co-occurrence within the Staphylococcal family (Figures 2B and 2C). The *sroB* gene, Aureowiki pan ID SAUPAN002490000, is annotated in 100% of the *S. aureus* strains in the Aureowiki metagenomic database. However, the *sroA* gene, Aureowiki pan ID SAUPAN002491000, is annotated in 88% of 33 strains within the Aureowiki metagenomic database (https://aureowiki.med.uni-greifswald.de/SAUPAN002491000). The *sroA* gene annotation is absent in the genome sequence maps of strains NCTC8325, Newman, RF122, and TCH60. We therefore decided to confirm that the *sroA* gene encoded a protein in *S. aureus* SH1000. We constructed a xylose-inducible FLAG-tagged allele of *sroA* (*sroA*-FLAG). We also constructed a xylose-inducible FLAG-tagged allele of *sroA* with a STOP codon substitution mutation for the start codon (*sroA*-FLAG^STOP^). We then performed western blot analysis using an antibody to the FLAG-tag on total protein isolated from xylose-induced exponentially growing cultures of SH1000 containing either 1.) empty vector control 2.) wild type *sroA*-FLAG, or 3.) *sroA*- FLAG^STOP^ (Figure 2D). We were able to detect SroA-FLAG protein levels from the wild type *sroA*- FLAG allele. However, expression from the *sroA*-FLAG^STOP^ allele was undetectable suggesting that the *sroA* gene does produce a protein (Figure 2D).

**Figure 2.**
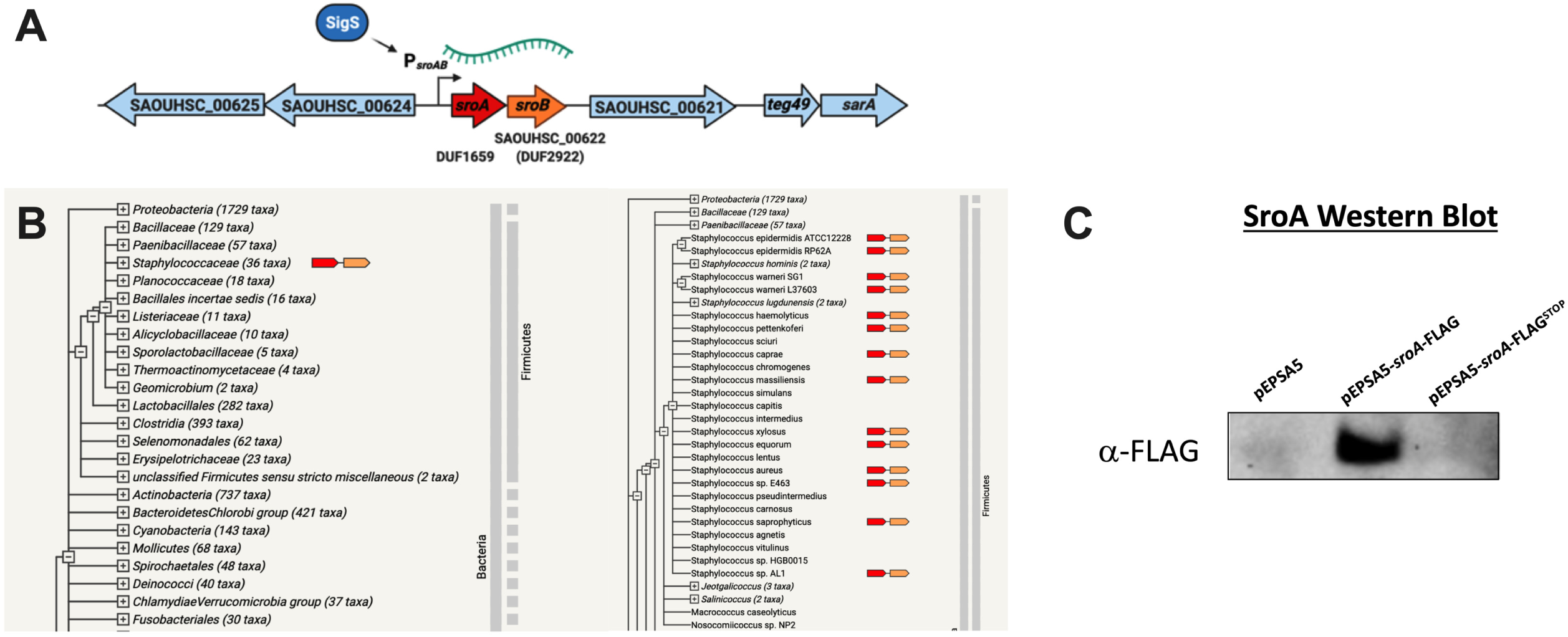
The *sroAB* genetic locus and co-occurrence within the *Staphylococceae*. A.) The *sroAB* genetic locus in *S. aureus* SH1000. B.) The amino acid sequence of SroA was translated from the *S. aureus* NCTC8325 genome and entered into the String database (string-db.org) and analyzed for co-occurance within bacteria. C.) FLAG-tagged alleles of SroA, wild type allele and allele with the start codon replaced with a stop codon (STOP), was over-expressed in log-phase growing *S. aureus* SH1000 from a plasmid- based xylose inducible promoter. Then total protein was isolated and analyzed by western blot using antibody to FLAG.

### SigS directly regulates the transcription of *sroAB* in response to DNA damage

To confirm that SigS over-expression induced the up regulation of *sroAB*, we conducted *sroAB* northern blot analysis on total RNA samples isolated from *S. aureu*s following SigS over-expression (Figure 3A). In the strain containing the vector control, both SigS and SroAB expression were completely undetectable. Conversely, in the strain containing the episomal xylose-inducible allele of SigS, SroA was strongly induced along with SigS. We previously reported that SigS activity is induced following exposure to the DNA damaging agent MMS (11, 13). We therefore reasoned that *sroAB* expression should also be induced by MMS in a SigS dependent manner if it is a regulatory target of SigS. To test this hypothesis, we grew wild type and Δ*sigS* strains of *S. aureus* HOU (MRSA) in TSB supplemented with MMS and analyzed total RNA by northern blot analysis. Consistent with our previous results, the *sigS* transcript was induced in cultures treated with MMS (Figure 4). The *sroA* mRNA was also detected in cultures treated with MMS, however, in the absence of *sigS*, the *sroA* transcript was undetectable. This result further supports the idea that SroA is a regulatory target of SigS. To determine if SigS directly binds to the *sroAB* promoter, we performed in vitro transcription using RNA polymerase Core enzyme complexed with purified recombinant SigS protein and analyzed transcriptional yields using endpoint RT-PCR. RNAP core reconstituted with purified recombinant SigS did indeed result in a transcription product, suggesting that the *sroAB* promoter region is recognized by the RNAP-σ^S^ complex and that *sroAB* is a direct target of SigS.

**Figure 3.**
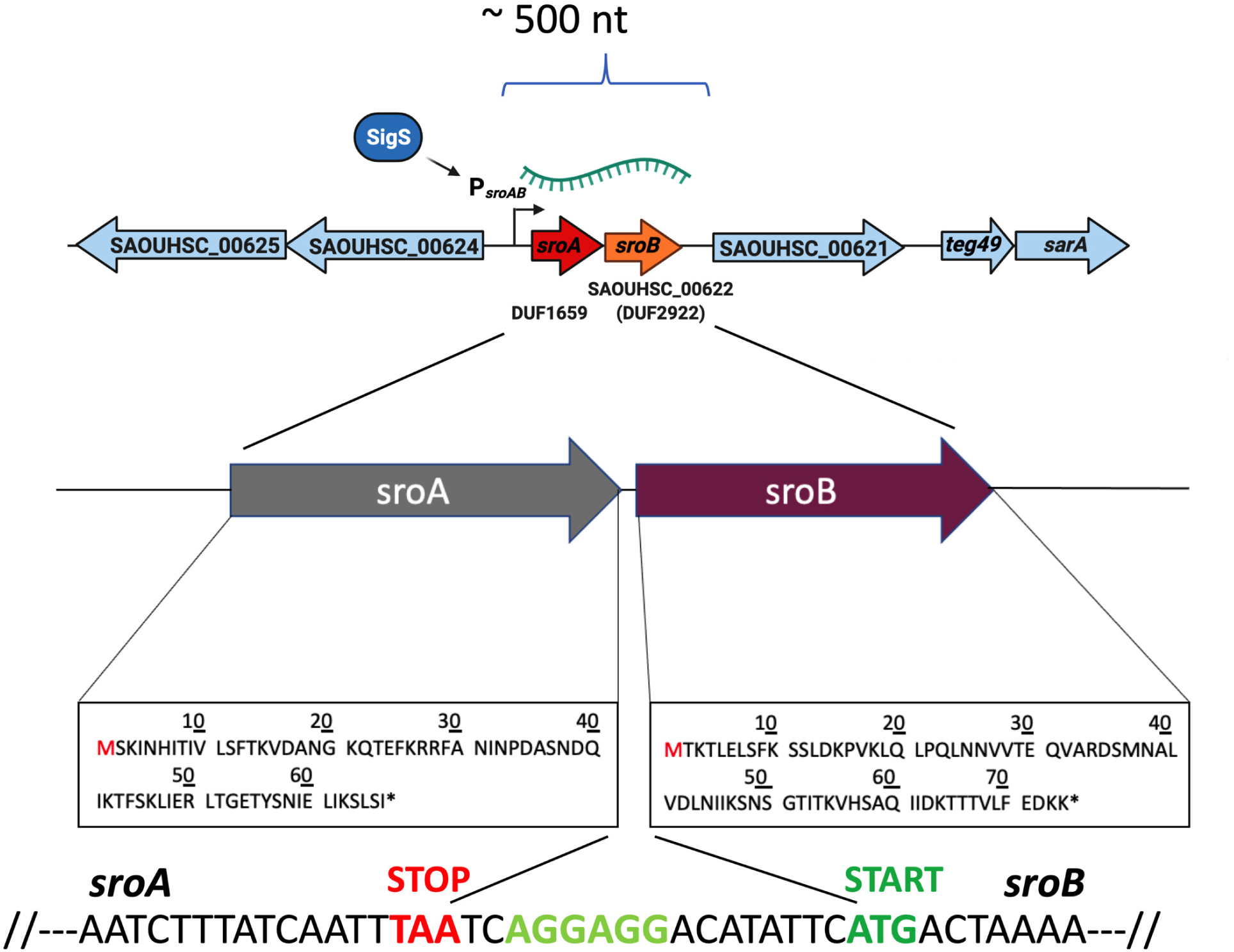
The *sroAB* locus, SroAB protein sequences, and Shine Dalgarno sequence. The *sroAB* genetic locus with the amino acid sequence for each gene magnified. The sequence between the *sroA* STOP codon (red letters) and the *sroB* START codon (dark green letters) highlighting a canonical shine-delgarno (ribosome binding site) sequence between them (lime green letters).

**Figure 4.**
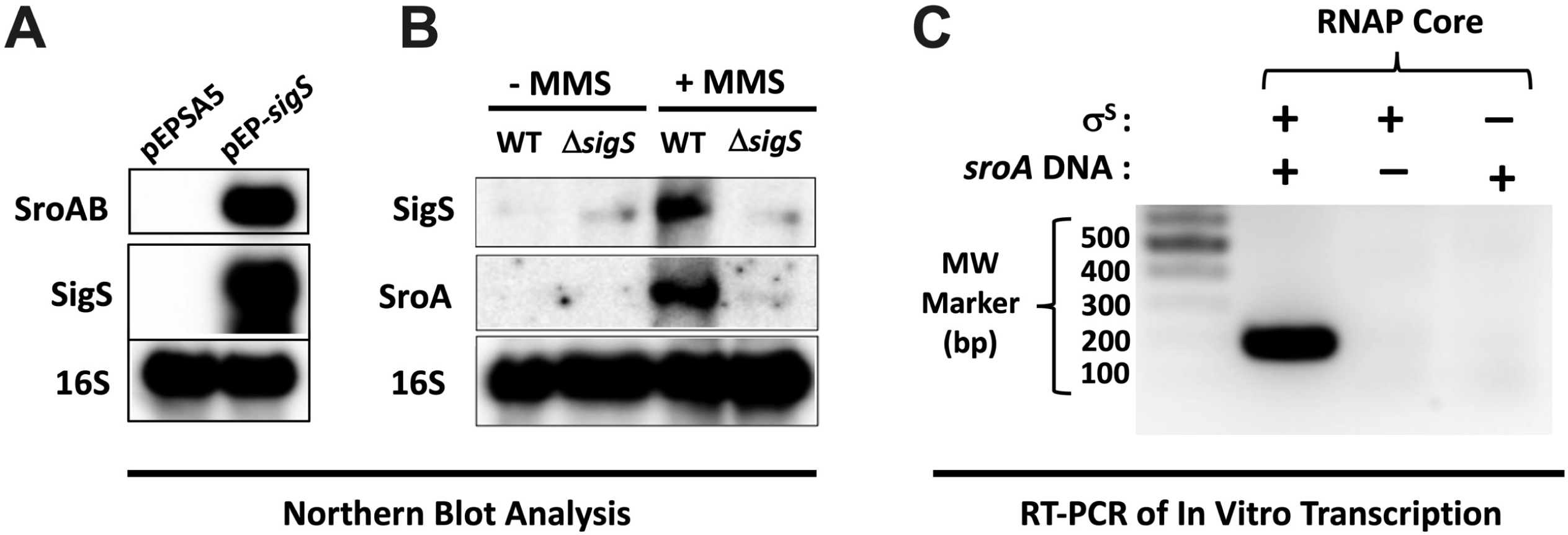
Northern Blot and In vitro transcription of *sroAB* in response SigS activity. A.) Overnight cultures of wild type SH1000 carrying pEPSA5 or pEPSA5-*sigS* were sub cultured (1:200) in 30 mL of TSB-Cm and grown to an OD_600_ of 0.3. Cultures were induced with xylose, to a final concentration of 2%, for 30 minutes and total RNA was isolated. Total RNA (3 μg) was analyzed by Northern Blot using Biotin labeled probes annealing to for 16S rRNA, SigS mRNA, and SroA mRNAs. B.) Overnight cultures of *S. aureus* HOU wild type and 11.*sigS* were sub cultured (1:200) in 30 mL of TSB supplemented with MMS to a final concentration of 30 mM and grown to an OD_600_ of 0.3. Total RNA was analyzed by Northern Blot using biotinylated probes corresponding to 16S rRNA, SigS mRNA, and SroA mRNA. C.)

### SroA and SroB have divergent effects on *sigS* mRNA and its antisense RNA transcript expression

Due to the intense induction of the *sroAB* transcript in response to SigS over-expression, we hypothesized that SroAB may exhibit feedback regulation on SigS expression as seen in other bacteria as it pertains to regulatory circuits involving alternative sigma factors (17). We decided to measure the expression of SigS mRNA upon over-expression of either *sroA*, *sroB*, or *sroAB* and measured both the *sigS* mRNA and the previously described *sigS* cis-antisense RNA (14). The *sigS* operon is hypothesized to be a post-transcriptionally repressed by s750, which is encoded directly downstream of SAOUHSC_1899 (SACOL1829) and has an extended 3’ UTR (s752) that spans the *sigS* operon. Therefore, we decided to also measure the accumulation of the s750 (SigS cis-antisense RNA) in this experiment. Remarkably, over-expression of SroA induces *sigS* mRNA accumulation, while SroB over-expression induces accumulation of s750 (*sigS* cis-antisense RNA). Interestingly over-expression of the complete *sroAB* transcript did not induce expression of either the *sigS* or its antisense transcripts (Figure 4A). While this result was interesting, the molecular mechanism that could explain this was unclear.

To explain this observation, we hypothesized that SroA and SroB may regulate each other through direct protein-protein interaction; thus, when both are expressed, they effectively cancel the activity of each other out. We decided to test this hypothesis by executing a Bacterial Two-Hybrid assay (27). We spotted colonies co-transformed with plasmids for encoding pEB355-*sroA* and pEB354-*sroB* in a 2x2 array, allowing us to test for potential SroA–SroB, SroA-SroA, and SroB-SroB interactions (Figure 4B). Our positive and negative controls exhibited the expected Lac^+^ and Lac^−^ phenotypes, respectively (Figure 4B). The SroA-SroA and SroB-SroB groups were both Lac^−^ (Figure 4B). However, the SroA-SroB spots both exhibited a Lac+ phenotype (Figure 4B). Taken together, this suggests that SroA and SroB interact with each other, while not interaction with themselves.

### SroA inhibits SigS mRNA decay

While our data suggested that SroA promotes accumulation of *sigS* mRNA levels in the absence of an interaction with SroB, it was not clear how SroA promotes SigS accumulation. Since SroA lacks a predicted DNA binding domain, we hypothesized that the SroA stimulatory effect on SigS is either through an indirect stimulatory effect on *sigS* transcription or a direct effect on *sigS* mRNA turnover. We thus decided to measure *sigS* mRNA decay following MMS induction using a Rifampicin Chase Assay (Figure 5A). The *t*_1/2_ of *sigS* mRNA from *S. aureus* SH1000 containing the vector control was less than 5 minutes, while the *t*_1/2_ of *sigS* mRNA isolated from *S. aureus* SH1000 over-expressing SroA was approximately 15 minutes. This suggests that SroA acts to inhibit *sigS* mRNA decay (Figure 5A).

**Figure 5.**
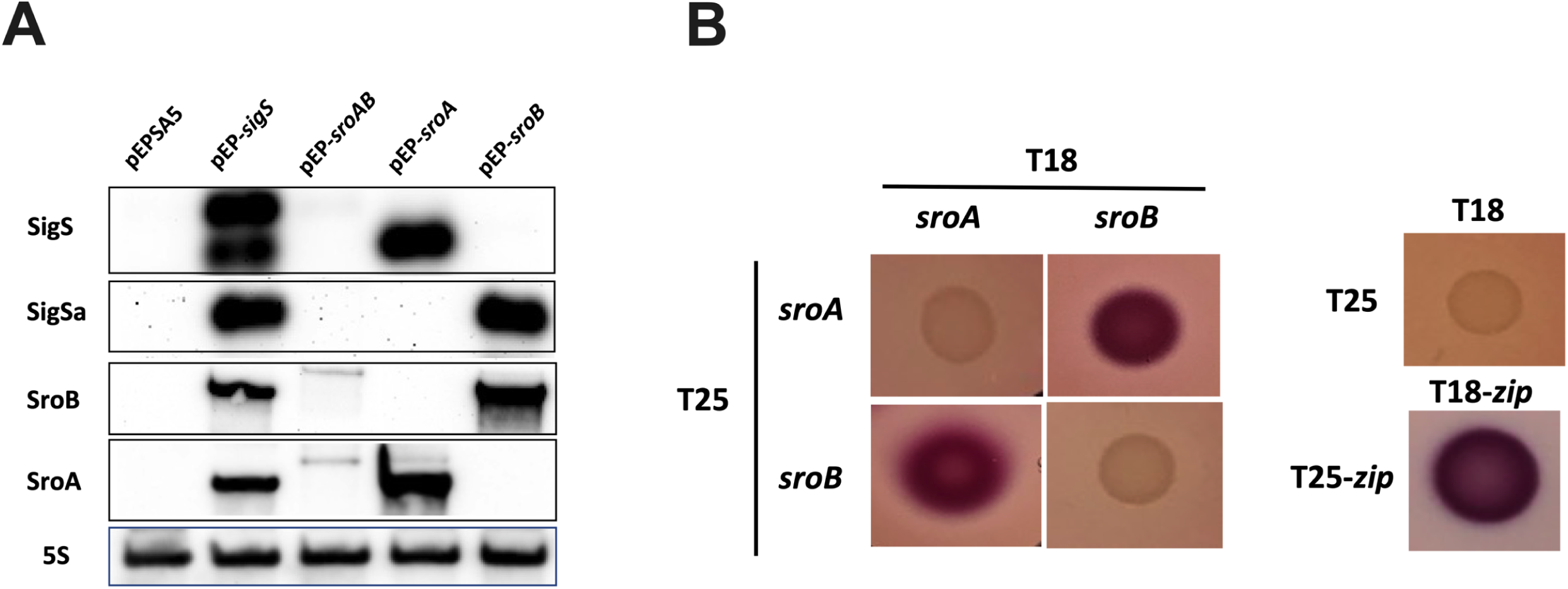
SigS and SigS antisense northern blot analysis and SroAB activity. A.) Over-night cultures of SH1000 carrying pEPSA5, pEP-*sigS*, pEP-*sroAB*, pEP-*sroA*, or pEP-*sroB* were sub-cultured in 30 mL of TSB-Cm and grown to an OD_600_ of 0.3. Cultures were induced with xylose, to a final concentration of 2%, for 30 minutes and total RNA was isolated. Total RNA (3 μg) was analyzed by Northern Blot using Biotin labeled probes annealing to for 16S rRNA, SigS mRNA, SigS antisense transcript, SroA mRNA, and SroB mRNA. B.) Bacterial Two Hybrid (BACTH) Analysis of SroA and SroB. The sroA and sroB were cloned into BACTH plasmids pEB355 and pEB354 and analyzed on MacConkey-Maltose agar plates at 30°C along with empty vector controls and positive controls (pEB355-*zip* and pEB354-*zip*). C.) Quantitative analysis of Bacterial Two Hybrid Analysis using β-galactose assay.

**Figure 6.**
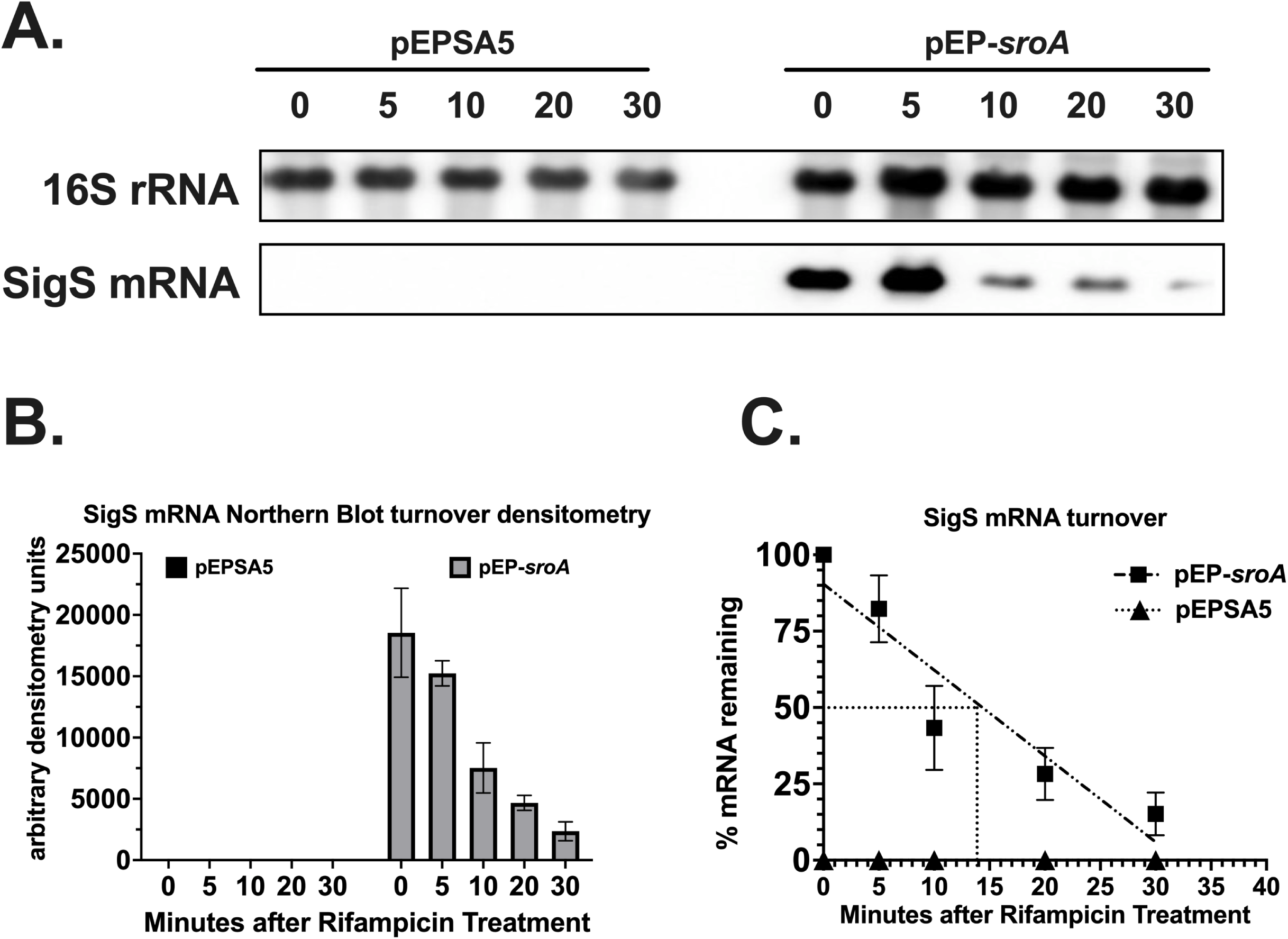
SigS mRNA turnover following over-expression of SroA. A.) Northern blot analysis was executed on wild type SH1000 carrying either pEPSA5 or pEP-*sroA* were grown in 30 mL of TSB-Cm supplemented with MMS to a final concentration of 25 mM to an OD_600_ of 0.3. Cultures were the induced with xylose, to a final concentration of 2%, and incubated for an additional 30 minutes. Cells were harvested and resuspended in fresh TSB-Cm supplemented with Rifampicin to a final concentration of 500 µg / mL and total RNA was isolated at 5 to 15-minute intervals. Total RNA (3 μg) was analyzed by Northern Blot using Biotin labeled probes annealing to for 16S rRNA, SigS mRNA, SigS and SroA mRNA. B.) Densitometry analysis of Northern analysis. C.) Calculation of the half-life (t*_1/2_*) of SigS based on densitometry analysis in section B. Quantitative Northern Blot data from Section B was subjected to linear regression analysis using GraphPad Prism 9. Experiments were repeated at least three times and data is presented as the mean plus or minus the standard error of the mean.

**Figure 7.**
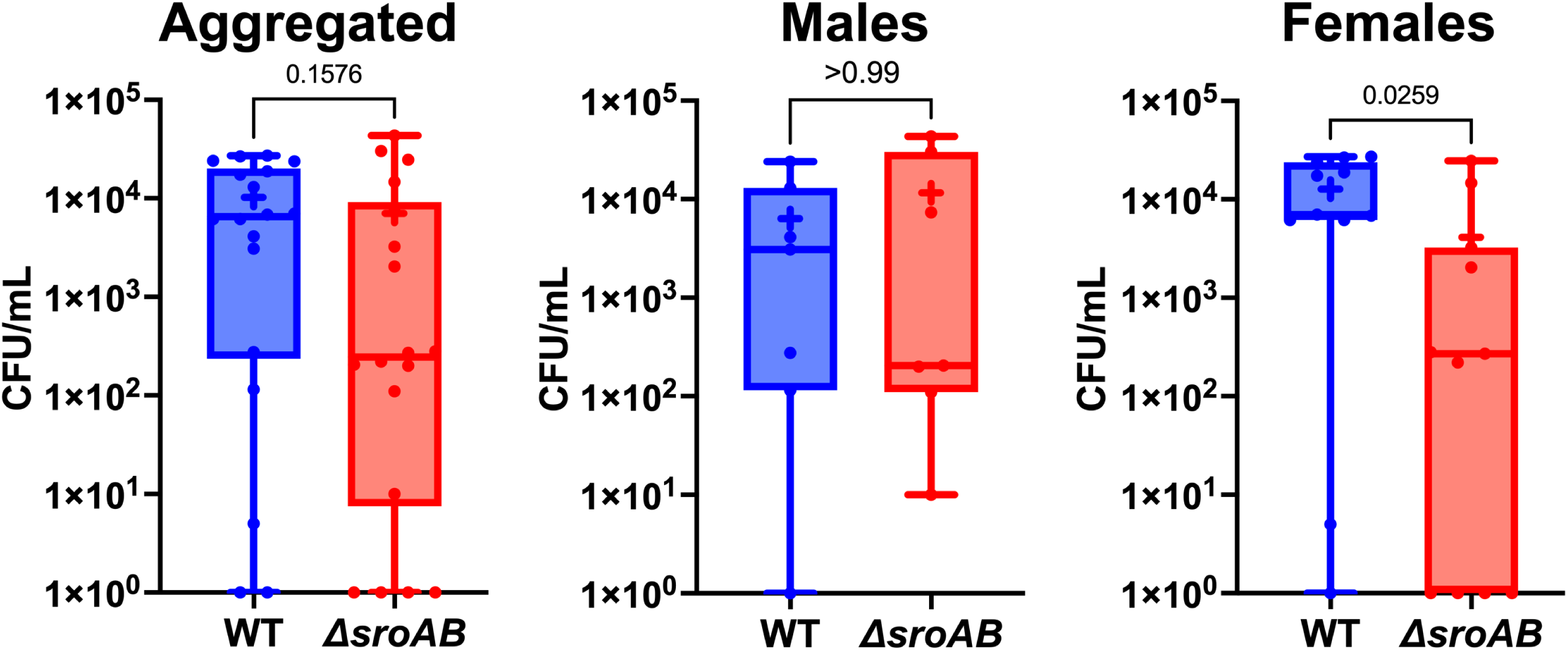
*S. aureus* virulence in a mouse model of pneumonia in the absence of *sroAB*. C57BL/6J mice (7 male, 11 female) were each infected intranasally with 5 x 10^7^ CFU of either USA300 (WT) or Δ*sroAB*. After 24h, mice were sacrificed, lungs harvested, and bacterial burden determined. Shown are max and min values (whiskers), and 25-75 percentile (boxes). Horizontal lines = mean; + = median. Statistical significance was determined using a Mann-Whitney test.

### The *sroAB* operon is required for full virulence in female mice

While SroA clearly acts to regulate SigS abundance at the post-transcriptional level, the role of the *sroAB* locus in virulence is undefined. We hypothesized that the *sroAB* locus is necessary for *S. aureus* virulence since SigS promotes virulence, and SroA stimulates *sigS* expression. To test this hypothesis, we measured virulence of a *sroAB* double mutant in a murine model of pneumonia. C57BL/6J mice were each infected intranasally with wild type and Δ*sroAB* strains of *S. aureus* USA300 (MRSA strain). After 24h, mice were sacrificed, lungs harvested, and bacterial burden determined as colony forming units per ml. Upon analysis we noted a statistically significant, threefold decrease in the amount Δ*sroAB* mutants recovered from female mice compared to wild type. Interestingly, there was no difference in the amount of wild type or mutant cells recovered from male mice, for reasons that are not clear.

## DISCUSSION

Through our SigS transcriptome studies, we identified a previously uncharacterized locus encoding a bicistronic transcript, that we named *sroAB*, induced 1000-fold following SigS over-expression. The *sroAB* transcript was the most highly induced transcript upon transient over-expression of SigS. This suggests that SroA and SroB are major downstream effectors of the SigS stress response. SroA and SroB each encode mid-sized bacterial proteins with 68 and 72 amino acid residues, respectively. While these proteins are just above the upper limit of 50 amino acids for the definition of short ORFs in bacteria, their relatively small size is interesting to note as investigations into the dark proteome and dark genome are increasing (30–33).

We also identified several immune evasion genes that were up regulated in response to SigS over-expression (Figure 1C). Both Spa and Sbi are immunoglobulin binding proteins that would ultimately inhibit antibody opsonization to, phagocytosis of, and clearing of *S. aureus* infection. The other SigS up-regulated immune genes, *hlgBC* and *lukAB*, encode leukotoxins that are effective in the targeting of various white blood cells. Both operons are also stimulated by the SaeRS two component signal transduction system that sense human neutrophil peptides (ref). SigS-dependent up-regulation of *sbi*, *spa*, and *hlgB*C may provide a mechanism to explain the increased sensitivity of 11.*sigS* mutants to whole human blood and macrophages (13). Interestingly, many of the SigS repressed genes were a part of the SigB and CodY regulons. Repression of SigB activated genes upon over-expression of SigS could be due to sigma factor competition (SigS vs SigB) for core RNA polymerase (34, 35). Yet, the influence of Sigma factor competition in firmicutes is not clear.

A major function that we uncovered was the ability of SroA promote positive feedback regulation the of SigS. Previous studies have demonstrated that in vitro expression of SigS is very low. Several chemicals stimulate transcription of SigS, most notably the chemical mutagen MMS (11, 13). Following MMS treatment, *sigS* transcription in strongly induced. However, the mechanisms used to tightly control SigS expression and transduce chemical stressors to stimulate gene expression is still unclear. Biochemical and genetic screens for SigS transcriptional regulators yielded several candidates, some of which bind to the *sigS* promoters, to include KdpE and CymR (11, 13, 14). While ArlR and LacR are indirect regulators of *sigS* transcription (11, 13, 14). Tight regulation of *sigS* transcription is likely to be part of the mechanism used by S. aureus to repress levels and conserve cellular resources until SigS is needed for restoration of homeostasis after exposure to various stressors. However, transcriptional control is only likely to be part of the collection of regulatory switches needed to tightly control SigS levels.

Post-transcriptional control of gene expression usually includes small RNA regulation of translation or mRNA turnover. Prior to this work, little was known about the post-transcriptional regulatory control of SigS. Interestingly, we demonstrate a role for the *sroAB* locus in post-transcriptional regulatory control of SigS. We identified a role for SroA in the stabilization of the SigS mRNA transcript. SroA stabilization of the SigS mRNA is particularly interesting, given the intrinsically low levels of SigS and incomplete picture of SigS regulatory control. In addition to tight transcriptional control, low levels of endogenous SigS expression could be due to SigS mRNA instability; and SroA activity could mitigate this instability to transiently increase SigS levels. Interestingly, over-expression of SroB, nor SroAB together, did not have the same results as SroA over- expression. A positive Bacterial Two Hybrid interaction between SroA and SroB suggest that an SroA-SroB interaction inhibits the ability of SroA to stabilize SigS mRNA.

We created a model for a SigS-SroAB regulatory circuit in *S. aureus* (Figure 8). In the present of DNA damage, SigS transcription is induced. However, the level of SigS protein that is synthesized is relatively low due to high SigS mRNA turnover. The SigS protein that is synthesized directly stimulates transcriptional of the *sroAB* locus, leading to the expression of SroA and SroB. While SroA can stimulate SigS accumulation at the level of mRNA stability, a direct interaction SroA and SroB inhibits this. Under a condition that has yet to be identified, either 1.) The SroA–SroB interaction is inhibited (perhaps through a post-translational modification) or 2.) SroB is degraded by an intracellular protease. We postulate that free SroA acts promote SigS mRNA stability by inhibiting one or more RNase that promotes SigS turnover. The precise role of RNases in SigS mRNA turnover and the molecular mechanisms that control an SroA-SroB interaction are the subject of ongoing investigations in our laboratory.

**Figure 8.**
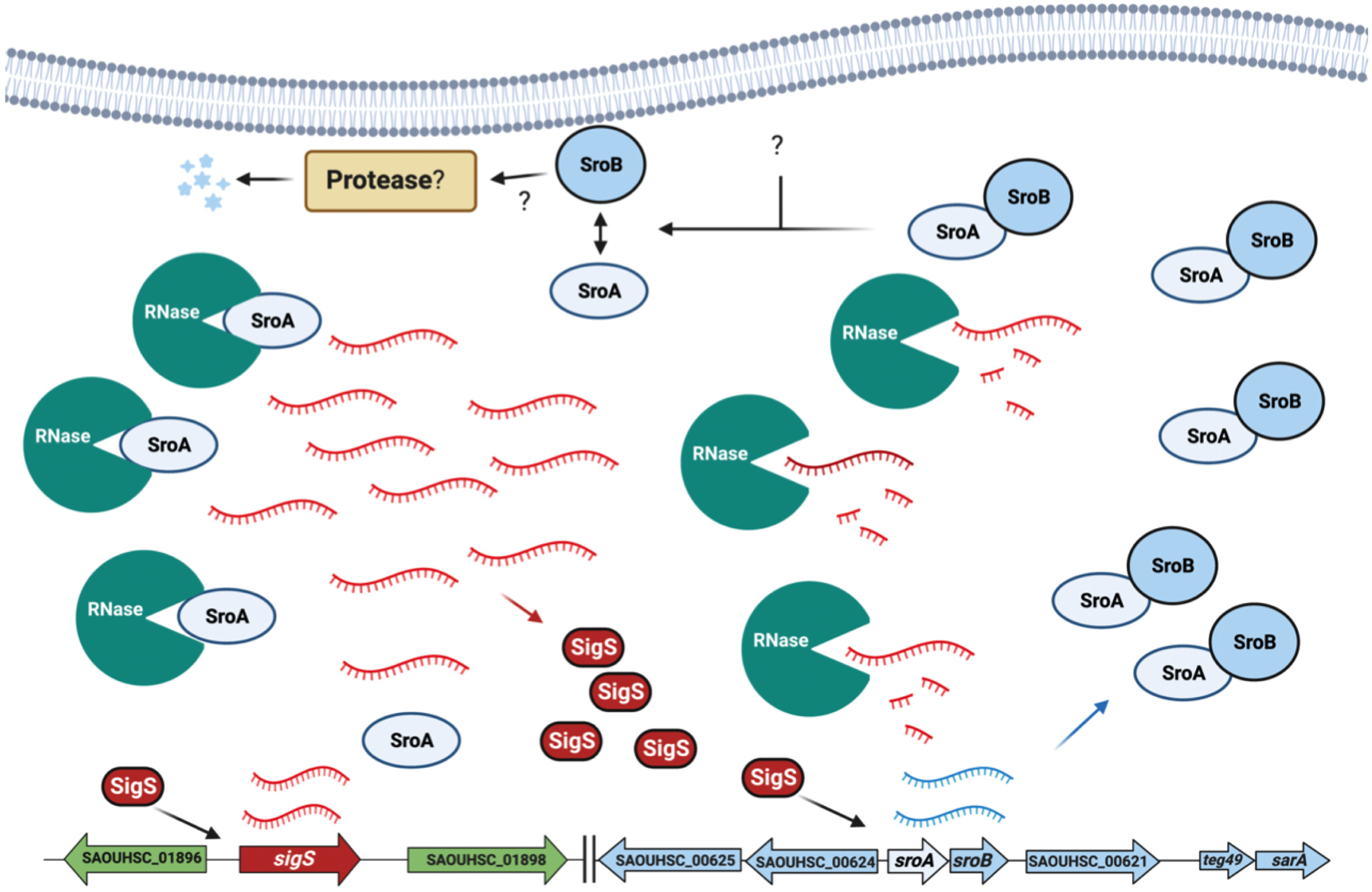
Model for The SigS – SroAB regulatory Circuit. In response to DNA damaging agent MMS, SigS is activated (through an unknown mechanism). SigS acts stimulate *sroAB* expression. SroA and SroB interact with each other under normal circumstances. In response to an unidentified internal or external signal, SroB is either inactivated or degraded. Then, free SroA promotes SigS accumulation through inhibition of RNases that promote SigS transcript degradation.

Prior to this work, it was well established that SigS was necessary for *S. aureus* survival in the presence of chemical stressors, environmental stressors, and exposure to immune cells. Specifically, these stressors include ethanol, H_2_O_2_, cell wall targeting antibiotics, chemical mutagens (MMS), elevated temperature, and intracellular macrophage survival. We aimed to identify downstream effectors of the SigS stress response to further understand SigS function and *S. aureus* infection. Our work supports a role for the *sroAB* locus in *S. aureus* virulence, and the need for further studies of SroAB function, which will increase our understanding of *S. aureus* virulence, and SigS mediated pathogenesis, immune evasion, and stress response; all of which are ongoing studies in our lab.

## Acknowledgements

This study was supported (in part) by several funding sources. These include a grant from the National Institute on Minority Health and Health Disparities of the National Institutes of Health under Award Number 2U54MD007597; grants AI124458 and AI157506 (L.N.S) from the National Institutes of Allergy and Infectious Diseases; a grant from The Bridge Fund and Pilot Project Program (K.M.T) of Howard University College of Medicine, and finally a grant from The Founders Chair in Basic Science from The Howard University Medical Alumni Association (K.M.T). We would like to thank members of the Thompson and Shaw Labs for critical review of this manuscript.

